# A Scalable High-Density Microwell Assay for Single-Cell Clonal Expansion Profiling

**DOI:** 10.64898/2026.04.10.717842

**Authors:** Stefanius Karoliina, Raut Shiska, Presley Bryan, Digant P Dave

## Abstract

Traditional clonogenic assays remain central to evaluating the self-renewal capacity of tumor cells. However, the assay relies on subjective endpoint measurements, is restricted to two-dimensional monolayer growth, and lacks the single cell resolution required to resolve heterogeneous expansion behaviors. We describe a high-density microwell array-based platform for quantitative assessment of single cell clonogenic growth outcomes, defined by cell count distributions spanning non-dividing, slow-dividing, and fast-dividing three-dimensional colony forming phenotypes. This approach links initial single-cell occupancy to defined growth outcomes across thousands of indexed microwells per well. The platform integrates high-density, low-adhesion microwell arrays within industry standard device plate formats and an automated image analysis pipeline incorporating machine learning, enabling parallel quantification of spatially indexed founder-derived microwells using widely accessible automated imaging systems. The assay was implemented in both 4-well and 96-well plate formats to evaluate reproducibility and scalability across different plate configurations. Using three glioblastoma cell lines as model systems, we demonstrate reproducible single founder occupancy and consistent clonal growth outcome distributions across replicate formats. This integrated microscale assay platform enables systematic quantitative characterization of clonogenic expansion capacity at single cell resolution and is compatible with applications in cancer biology, therapeutic testing, and functional single cell phenotyping. By resolving single-cell persistence, limited expansion and high expansion outcomes within a scalable high-density format, this approach expands the analytical resolution of single cell clonogenic profiling beyond traditional binary colony scoring.

## 1. Introduction

Normal somatic cells exhibit a controlled and finite replicative capacity, whereas the ability to generate unlimited progeny is associated with tumor initiating potential and stem cell-like behavior.[1-3] The clonogenic assay, first described in the 1950s by Puck and Marcus, is a benchmark *in vitro* method for assessing clonogenic potential and self-renewing capacity of tumor cells.[1, 4, 5] It is widely regarded as the gold standard for evaluating cytotoxic effects, particularly in tumor therapy response studies.

Despite its longstanding role as the gold standard colony formation method, the classical clonogenic assay has fundamental limitations that constrain its scalability and relevance for modern screening needs. Traditionally conducted in low-throughput 6- or 12-well plate formats, it is poorly suited for evaluating combination therapies or multiple drug concentrations in parallel, limiting the number of therapeutic conditions that can be interrogated within a single experiment.[4, 6] The workflow remains highly manual, operator-dependent, and sensitive to subtle variations in seeding and handling, contributing to inter-laboratory inconsistency.[5-8] Analysis typically relies on single endpoint colony counting, an inherently subjective process further complicated by the arbitrary convention of defining a colony as ≥50 cells or a diameter of ≥50 microns.[4, 5] Because classical clonogenic assays evaluate only the final colony outcome, they lack the dynamics leading to colony formation at single cell resolution, and therefore cannot link endpoint colony size to the initial founder cell. Furthermore, traditional clonogenic assays are performed on large two-dimensional monolayer surfaces where colonies expand laterally, frequently overlap, and cannot be segmented or tracked automatically, limiting both quantification and automation. Additionally, the assay does not recapitulate 3D clonal growth that is seen *in vivo*.[4] Although recent improvements, including plate-based miniaturization, automated imaging, and machine learning-based colony detection have helped streamline analysis [5, 9-12], these adaptations still depend on endpoint colony scoring and typically apply fixed size threholds. As a result, intermediate expansion states and single-cell persistence outcomes are often collapsed into a single non-colony category, limiting resolution of differences in growth capacity among tumor cells within the same population.

In response to these limitations, several adaptations have sought to modernize clonogenic workflows. For example, microstructured and 3D culture systems, such as collagen-based matrices, hydrogel-supported growth, and microwell platforms, have improved environmental control and enabled live-cell observation of early clonal events. [13-17] These developments represent important progress but still rely on population-level or endpoint-based measurements, offering limited ability to preserve outcome information linked with the initial founder cell state at large sampling scale. Moreover, many microengineered platforms require custom microfluidics, and specialized imaging setups limiting their accessibility and preventing widespread adoption of single cell resolved clonogenic analyses.[16] Even when confined microenvironments are available, most systems cannot couple that confinement with scalable imaging and automated analysis, leaving a gap between biological relevance and practical usability.

We have developed a clonogenic assay platform for quantitative assessment of single cell– derived clonal growth outcomes at large sampling scale, spanning single-cell persistence, limited expansion, and high expansion three-dimensional colony forming phenotypes. Unlike classical approaches that quantify clonogenicity as a single endpoint measure, our methodology preserves distribution-level information and provides a more detailed view of differences in clonal growth capacity within a tumor cell population. To implement our methodology at a high-throughput scale, the design and manufacturing of ultra-high density microwell array plates, and the development of an automated image-analysis workflow that links endpoint proliferation outcomes to the initial founder state of each indexed microwell, were required. The approach integrates hardware design, automated imaging compatibility, and computational analysis to enable large-scale quantification of clonal growth outcomes across tens of thousands of indexed microwells at single cell resolution. The high-density microwell platform supports spatial indexing of individual founder cells and quantitative endpoint analysis across tens of thousands of clonal units.

Our engineered microwell array plates in standard well format layouts, consist of ∼10,000 microwells per macrowell, each with uniform 50 µm × 50 µm × 50 µm geometry patterned onto a cyclic olefin polymer (COP) film. Importantly, this integrated microwell design achieves experimental scales that would otherwise require large numbers of standard multiwell plates when using single cell dispensing approaches for clonal assays. Conventional 96-, 384-formats can isolate single cells, but reaching the same number of clonal units as a single microwell array plate demands orders of magnitude more plates and associated single cell dispensing instrumentation, dramatically increasing reagent usage, labor, imaging burden, and total assay time. Such distributed plate-based workflows also introduce additional sources of technical variability, reducing consistency across replicates. In contrast, our microwell array platform consolidates thousands of uniformly structured clonal units into a single plate, offering far greater throughput, tighter control of culture conditions, and more reproducible quantification of single-cell clonal growth outcomes. Plates are produced in both 4-well and 96-well formats, maintaining compatibility with standard automated imaging systems, while significantly reducing reagent consumption and incubator space requirements.

The microwell substrate is coated with Polyethylene glycol (PEG) to minimize cell adhesion and promote cellular confinement with defined microwell boundaries, which is important for accurate quantification of clonal growth capacity under non-adherent conditions. [18, 19] PEG coating is widely used in 3D tumor spheroid assays as a low adhesion coating. [20] The microwell architecture organizes cells into discrete spatially indexed positions, improving clonal separation and simplifying automating machine learning (ML)-based segmentation and tracking, while also maintaining compatibility with standard automated imaging systems.

To test and validate our clonogenic assay platform, we chose Glioblastoma (GBM) tumor model which represents an especially relevant context for evaluating clonal behavior[21-23] because it exhibits pronounced intratumoral heterogeneity.[24-26] Rare slow-cycling or quiescent GBM cells contribute to residual disease and post-treatment regrowth, however, such behaviors are not resolved by traditional colony-formation assays. [27-29] To reflect this biological diversity, we selected three widely used glioma cell lines with well-documented differences in clonogenic potential and growth behavior: U251, a robust and frequently used line in clonogenic assays [4, 5]; U87MG, characterized by limited self-renewal [30, 31]; and T98G, a similarly low-clonogenic and less-proliferative line with distinct proliferative constraint. [14, 32-34] Together, these models span a representative spectrum of glioma growth phenotypes and provide a rigorous testbed for evaluating the reproducibility and generalizability of the assay.

## 2. Materials and Methods

### 2.1 Clonogenic Microwell Array Plate Design and Fabrication

Our microwell array plate consists of a microfabricated bottom film (Fig. S2A) bonded to a top reservoir frame using a biocompatible pressure-sensitive adhesive (PSA) (Fig. 1A). Two plate layouts were produced: a 4-well format for pilot studies, which has a footprint of a standard glass slide, and a standard 96-well format. Each of these layouts is compatible with high-throughput imaging microscopes. Each microwell/reservoir contains approximately 10,000 microwells arranged in a grid (Fig. 1A–B), with a total of 40K and 960K microwells in 4-well and 96-well plates, respectively. Each microwell measures 50 × 50 × 50 µm (length × width × depth) with a 20 µm spacing between microwells (Fig. 1B, S2). These microwells provide well defined spatial confinement to support clonal outgrowth of single cells into three-dimensional colonies (Fig. 1C).

**Figure 1.**
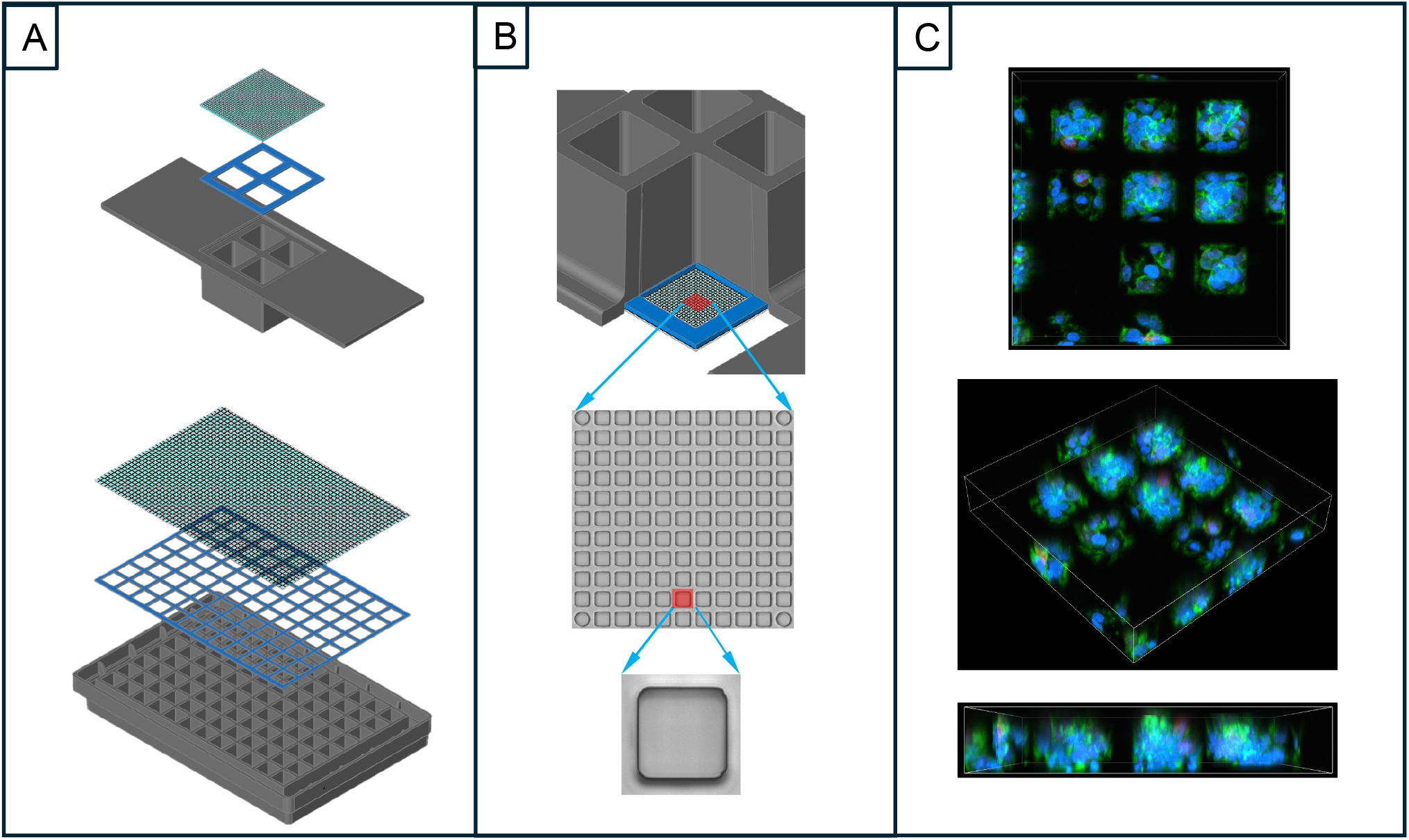
Design and assembly of the microwell array plate. (A) Exploded schematic illustrating the three components of the plate: a microfabricated bottom film containing the microwell array, a bottomless reservoir frame available in 4-well (top panel) and 96-well formats bottom panel), and a die-cut pressure-sensitive adhesive (PSA) tape layer used to bond the components (B) Each well contains ∼10,000 microwells, each measuring 50 × 50 × 50 µm. (C) Representative multichannel confocal 3D image stack showing clonal outgrowth of U251 cells in microwells. Panels depict an isometric view (middle), top view (top), and side view (bottom). Nuclei are shown in blue and F-actinin green.

**Figure 2.**
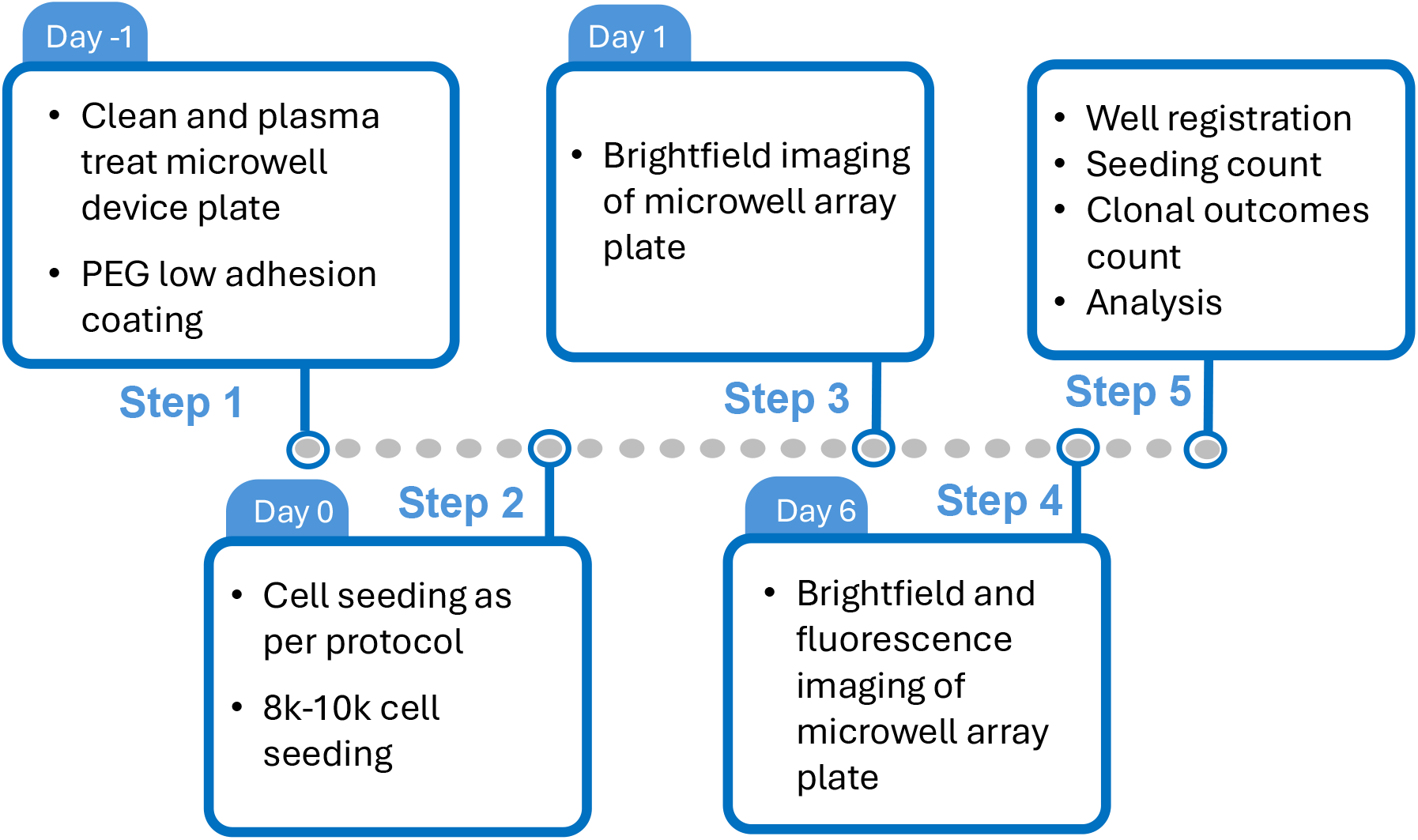
Workflow of sequential steps involved in conducting the assay. Step 1 (day -1) involves preparing the microwell plate for initiating the assay. Step 2 (day 0) cells are seeded. Step 3 (day 1) label-free bright field imaging of microwell array plate is performed. Step 4 (day 6) reimaging of the microwell array plate in bright field and fluorescence (nuclear stain with Hoechst dye) channels. Step 5 involves computational analysis of the data collected in Step 3 and Step 4.

**Figure 3.**
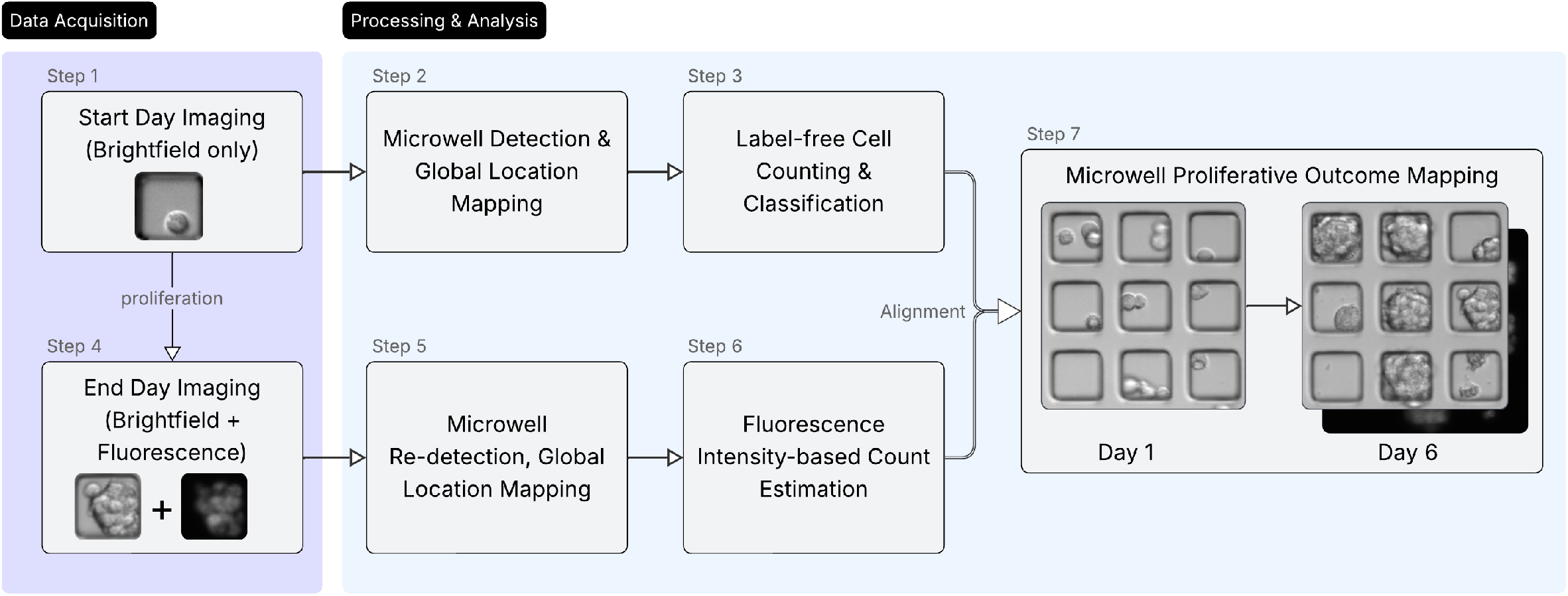
Image analysis pipeline for tracking clonal outcomes in microwell devices. Day 1 brightfield images are used for microwell detection, global location mapping, and label-free cell counting to classify each well at seeding. At the end of the experiment, brightfield and fluorescence images are re-acquired, microwells are re-detected and aligned to their Day 1 coordinates, and cell numbers are estimated from Hoechst intensity. Combining Day 1 classification with Day 6 counts enables microwell-level fate tracking for single-founder and multi-founder microwells. Day-1 and Day-6 composite images to classify founder occupancy and assign linked proliferation outcomes based on predefined cell-count categories.

Plate manufacturing was performed in three stages: (i) fabrication of the microwell array bottom film, (ii) fabrication of the top frame, and (iii) final assembly. The microfabricated bottom film containing the microwell array was printed on a 6″, 188 µm-thick cyclic olefin polymer (COP) substrate using a precision micro-thermoforming fabrication process (Supplementary Fig. S1A). Briefly, a 6” COP sheet was heated above its glass transition temperature and pressed against a microstructured transfer mold containing an array of negative microwell grid features. Under controlled temperature and pressure, the softened polymer conformed to the mold geometry, replicating the microwell structures with high fidelity (Supplementary Fig. S1B). After cooling below the glass transition temperature, the patterned COP sheet was released from the mold, yielding a mechanically stable, optically transparent microwell array suitable for cell culture and imaging applications. The printed microwell array shows high dimensional reproducibility and tolerance to design specifications.

The rationale for the choice of COP as the substrate for the bottom film is threefold: 1) It is highly compatible with scalable thermoplastic microfabrication molding technologies such as hot embossing and thermoforming to imprint reproducible microfeatures on thin films, 2) It is biocompatible for cell culture applications and does not absorb small drug molecules, which is critically important for drug studies, and 3) COP has superior optical properties in comparison to other widely used thermoplastics used in plateware such as polystyrene. Optical properties of COP are comparable to borosilicate glass used in coverslips for microscopy applications. COP has glass-like transparency in the visible (> 350 nm) and near-infrared spectrum of light and low birefringence. Optical thickness (physical thickness x refractive index) of microstructured COP film combined with its optical properties make it suitable for high-resolution brightfield and fluorescence imaging modes. The choice of film thickness (188 µm) is deliberate, to enable high-resolution confocal imaging and facile film removal for downstream analysis.

The bottomless top frame is manufactured by standard injection molding using black COP (black dye mixed with COP pellets). The black dye is used to facilitate better imaging quality due to reduced reflection and scattering of excitation light for fluorescence imaging. The injection molding process involves creating a negative mold cavity by precision machining hardened tool-steel and placing it into an injection mold system. The COP pellets are introduced into the injection molding system, where they are heated above the glass transition temperature and injected into the mold cavity under controlled conditions. Once injected into the cavity, the mold is cooled below the melting point, and the part is ejected from the tool to finish cooling.

The bottom film and PSA are die-cut to an appropriate dimensional fit for 4-well and 96-well plate format and bonded together under controlled pressure.

### 2.2 Reagents, Cell Lines and Culture Conditions

PLL-g-PEG (Nanosoft Biotechnology LLC) was prepared at a stock concentration of 1 mg/mL in phosphate-buffered saline (PBS; Thermo Fisher Scientific), aliquoted, and stored at −20 °C until use. Hoechst 33342 solution was purchased from VWR International, LLC.

U87MG and T98G glioblastoma cell lines were obtained from the American Type Culture Collection (ATCC). U251 cells were provided by Dr. Bachoo (UT Southwestern Medical Center). U87MG and T98G were maintained in Eagle’s Minimum Essential Medium (EMEM; ATCC) supplemented with 10% (v/v) fetal bovine serum (FBS, Gibco) and 1% penicillin–streptomycin (Thermo Fisher Scientific). U251 cells were maintained in EMEM supplemented with 10% (v/v) FBS, 1% penicillin–streptomycin, 1% MEM Non-Essential Amino Acids, and 1% sodium pyruvate (Thermo Fisher Scientific).

Cells were passaged at 70–90% confluence using 0.05% trypsin-EDTA (Thermo Fisher Scientific). After detachment, complete medium was added to neutralize trypsin, and the cell suspension was gently pipetted to disperse any remaining clumps. The suspension was then passed through a 40 µm cell strainer to obtain a uniform single-cell suspension. Cell concentration and viability were measured using a TC20™ Automated Cell Counter (Bio-Rad) with trypan blue exclusion. All experiments were performed using cells at passage numbers below 20. Cell lines were maintained at 37 °C in a humidified incubator with 5% CO_2_ and were routinely tested and confirmed negative for mycoplasma contamination.

### 2.3 Clonogenic Assay Setup and Seeding Procedure

The clonogenic assay described by Franken et al. (2006) was used as the foundational protocol, with modifications to enable single-cell seeding and longitudinal imaging in the microwell array format. Prior to coating, each well of the 4-well and 96-well plates was filled with deionized water, followed by air drying with pressurized air. Plates were then plasma treated at15W power, 50% duty cycle, with 20% oxygen level at 350 mTor pressure.(Tergeo Plasma Cleaner - Pie Scientific) to remove organic contaminants and render the microwell surface hydrophilic for wettability.

Following plasma treatment, wells were rinsed with sterile PBS and coated with 0.1 mg/mL PLL-g-PEG in PBS. The coating solution fully covered the microwell surface, and plates were incubated overnight at 4 °C. On the following day, plates were equilibrated at 37 °C for at least 15 minutes. The coating solution was aspirated, and wells were rinsed twice with PBS to remove unbound polymer; a third PBS rinse was left in place until seeding. Plates were UV-irradiated in a biosafety cabinet for at least 15 minutes before use.

Cells were detached with 0.05% trypsin-EDTA, neutralized with complete medium, gently pipetted to dissociate remaining clumps, and passed through a 40 µm cell strainer. Cell concentration and viability were determined, and suspensions were diluted to the desired seeding density.

For each cell line, four independent 4-well plates were prepared (12 total 4-well plates across three cell lines). In addition, two independent 96-well plates were prepared, with all three cell lines seeded in each plate. Before cell seeding, each well was filled with 100 µL of complete medium. U87MG and T98G cells were seeded at 8,000 cells per well, and U251 cells at 10,000 cells per well, in a total volume of 150 µL. Plates were placed on a 3-mm orbit shaker at 140 rpm for 45 seconds to promote even distribution of cells into microwells. Plates were transferred to a humidified 37 °C incubator and cultured overnight. Day 1 label-free imaging (brightfield mode) was performed within 18 hours of seeding to document initial microwell occupancy. Wells were then topped up with complete medium as needed, and cultures were maintained for 6 days, with medium refreshed every third day.

At the assay endpoint, Hoechst 33342 staining was performed under live-cell conditions. A 7 µM working solution was prepared in fresh medium. Growth medium was aspirated and replaced with the staining solution, ensuring full coverage of each well. Cells were incubated at 37 °C for 15 minutes in the dark. After staining, wells were washed once with PBS and replenished with fresh medium. Imaging was performed immediately using epifluorescence microscopy (excitation ∼350 nm, emission ∼461 nm). Live imaging was performed under standard incubation conditions, using a stage-top incubator.

### 2.4 Categorization of Microwell Proliferation Outcomes

Microwells were first classified at Day 1 based on initial occupancy. Microwells containing exactly one cell were designated as single-founder microwells, whereas microwells containing more than one cell were categorized as multi-founder microwells and analyzed separately.

At Day 6, outcomes were assigned based on the number of cells present within each indexed microwell: 0 cells (no cells; NC), 1 cell (single-cell persistence; SP), 2-7 cells (limited expansion; LE), or ≥8 cells (≥8-cell expansion). For multi-founder microwells, the same numeric categories were applied; however, a Day 6 count of one cell was interpreted as reduction to a single cell (RS) rather than persistence of a single cell.

The ≥8-cell threshold was used as an operational boundary to distinguish higher expansion from minimal or limited proliferation within a single assay window. Exact cell counts were retained for all microwells, and finer stratification of higher cell-count ranges can be performed as needed; however, four-category framework was selected to maintain analytical clarity and align with the objective of distinguishing persistence, limited expansion, and higher expansion states.

### 2.5 Data Acquisition & Analysis Workflow

Plates were imaged using two automated microscopes (Agilent Cytation 5 and Thermo Fisher EVOS M7000) at 10× magnification in brightfield and fluorescence modes. The analysis workflow is designed to remain agnostic to the imaging system, provided that minimum requirements for spatial resolution and bit depth (12- or 16-bit) were met. For each well, partially overlapping image tiles (approximately 20% overlap) were acquired and stitched using the native microscope software. The resulting whole well composite images were exported for downstream analysis. Imaging was performed at two discrete time points: Day 1 following seeding and Day 6 following the proliferation period. Day 1 imaging was conducted in brightfield to establish a label-free record of initial microwell occupancy, and Day 6 imaging included brightfield and Hoechst fluorescence channels. Imaging was performed under controlled temperature and CO_2_ conditions to maintain standard culture parameters. No intermediate time-lapse imaging was required for growth outcome classification.

Day 1 brightfield composites were used to detect and spatially index individual microwells. Microwell boundaries were identified based on the patterned array geometry, and each microwell was assigned a unique coordinate within a global reference frame spanning the entire well. Because plates were removed from the microscope between imaging sessions, spatial correspondence required precise realignment. Circular reference markers integrated into the microwell array design and visible in brightfield images were used to establish a consistent global reference frame. The reference markers are replicated at 10 microwell intervals (Fig. 1B). A two-stage registration process was implemented: global alignment corrected plate-level translational shifts within the composite image, and local refinement corrected smaller positional deviations arising from minor stitching variation.

Following automated microwell detection and indexing, individual cells were segmented within each microwell from Day 1 brightfield images without fluorescent labeling. Brightfield imaging was intentionally used at seeding to enable label-free detection and avoid introducing fluorescent dyes prior to the proliferation period. An open-source segmentation model(Cellpose-SAM)[35] served as the base architecture. As pretrained models are typically optimized for a wide range of imaging conditions, additional adaptation was required to achieve optimal performance on the microwell imaging system. The model was therefore fine-tuned using a curated dataset of manually annotated Day 1 brightfield images specific to this platform. To accelerate the creation of training data, the pretrained model was first used to assist with semi-automated annotation. The resulting annotated dataset was then used to retrain the network on microwell-specific images, improving segmentation performance for this imaging context. Segmentation outputs were used to determine cell counts per microwell and classify microwells at seeding as single-founder or multi-founder, establishing the initial founder category for downstream outcome classification. The optimized model achieved a false-positive rate of 3.5% and a false-negative rate of 2.5% on an independent validation set.

At Day 6, cell quantification was performed using Hoechst fluorescence rather than instance segmentation. Because microwell geometry constrains lateral expansion and promotes 3D vertical cell stacking as cell numbers increase; densely populated microwells frequently contain overlapping nuclei in epifluorescence images. Under these conditions, direct instance segmentation is unreliable for accurate cell counting. A fluorescence intensity-based estimation approach was therefore implemented. For each well, isolated Hoechst-stained nuclei were automatically identified using size, circularity, and aspect ratio thresholds optimized for the respective cell line. More than 300 isolated single nuclei per well were used to generate a well-specific reference distribution of single-cell fluorescence intensity following background subtraction, noise filtering, and removal of outliers. The mean intensity of this filtered single-cell population was defined as the reference single-cell signal for that well. Total Hoechst fluorescence within each microwell was then measured and divided by the corresponding single-cell reference intensity to estimate cell number. This well normalization approach enabled automated and consistent quantification across a broad range of cell densities within the confined microwell architecture.

Well level clonal growth outcomes were assigned by integrating the Day 1 founder classification with the corresponding Day 6 estimated cell counts for each indexed microwell. Because microwell positions were defined within a global coordinate system and preserved through alignment, each microwell could be deterministically matched across time points. Clonal growth outcome categories were defined based on initial occupancy and final cell number, enabling quantitative clonal outcome distributions without reliance on continuous time-lapse imaging.

### 2.6 Statistical Analysis

Statistical analyses were performed to quantify intra-plate repeatability and inter-plate reproducibility of seeding and proliferation metrics. The replication structure is described in Section 2.3. Briefly, within each 4-well plate, individual wells served as technical replicates (intra-plate), and independent plates served as biological replicates (inter-plate). In the 96-well format, individual wells functioned as technical replicates, and inter-plate variability was assessed for the 4-well plates only. In 96-well plates, wells on the periphery were excluded from the analysis to remove edge effect.[36]

The coefficient of variation (CV) was calculated to summarize repeatability across technical replicates within plates, calculated separately for each 4-well or 96-well plate. Tables 1-2 summarize the CV values for seeding distributions and single-founder microwell outcomes. The CV values for multi-founder microwell outcomes can be found in the supplement (Table S1) along with inter-plate CV values for microwell occupancy, founder distributions, and single-founder clonal outcomes of the 4-well plates (Table S2, S3).

**Table 1.** Intra-plate seeding coefficient of variation (CV, %) for microwell occupancy and founder distribution among occupied microwells. Each value represents the intra-plate CV calculated across *n* wells in a plate.

| 4-well Plate Intra-Plate CV (%) for Seeding |  |  |  |  |  |  |  |  |  |  |  |
| --- | --- | --- | --- | --- | --- | --- | --- | --- | --- | --- | --- |
| T98G ($n = 4$) | | | | U251 ($n = 4$) | | | | U87 ($n = 4$) | | | |
| PID | Occ. | SFM | MFM | PID | Occ. | SFM | MFM | PID | Occ. | SFM | MFM |
| 1 | 1.12 | 2.03 | 5.64 | 5 | 5.05 | 5.87 | 6.02 | 9 | 2.17 | 2.53 | 3.92 |
| 2 | 4.56 | 6.48 | 4.98 | 6 | 1.05 | 3.93 | 4.38 | 10 | 3.17 | 4.88 | 7.43 |
| 3 | 3.64 | 4.90 | 5.25 | 7 | 1.36 | 2.31 | 0.95 | 11 | 2.88 | 2.05 | 4.35 |
| 4 | 0.74 | 2.86 | 8.83 | 8 | 3.98 | 3.46 | 5.08 | 12 | 5.24 | 4.04 | 8.09 |

| 96-well Plate Intra-Plate CV (%) for Seeding |  |  |  |  |  |  |  |  |  |  |  |
| --- | --- | --- | --- | --- | --- | --- | --- | --- | --- | --- | --- |
| T98G ($n = 16$)* | | | | U251 ($n = 24$)* | | | | U87 ($n = 16$)* | | | |
| PID | Occ. | SFM | MFM | PID | Occ. | SFM | MFM | PID | Occ. | SFM | MFM |
| 1 | 7.21 | 8.54 | 7.87 | 1 | 5.53 | 4.28 | 9.50 | 1 | 6.83 | 6.46 | 8.68 |
| 2 | 7.92 | 6.78 | 11.06 | 2 | 6.51 | 7.28 | 6.67 | 2 | 7.70 | 6.98 | 10.36 |

PID - Plate ID; Occ. - microwell occupancy; SFM - single-founder microwells; MFM - multi-founder microwells. \* All three cell lines were seeded on the same 96-well plate; edge wells excluded from analysis; $n$ indicates the number of wells (technical replicates) for the cell line.

**Table 2.** Intra-plate coefficient of variation (CV, %) for Day 6 clonal growth outcomes for single-founder microwells across 4-well and 96-well formats. Each value represents the intra-plate CV calculated across *n* wells in a plate.

| 4-well Plate Intra-Plate CV (%) for Clonal Growth Outcomes (SFM) |  |  |  |  |  |  |  |  |  |  |  |  |  |  |
| --- | --- | --- | --- | --- | --- | --- | --- | --- | --- | --- | --- | --- | --- | --- |
| T98G ($n = 4$) | | | | | U251 ($n = 4$) | | | | | U87 ($n = 4$) | | | | |
| PID | NC | SP | LE | $\geq 8$ | PID | NC | SP | LE | $\geq 8$ | PID | NC | SP | LE | $\geq 8$ |
| 1 | 6.02 | 8.55 | 3.87 | 17.74 | 5 | 7.57 | 6.67 | 15.43 | 2.73 | 9 | 3.24 | 11.08 | 2.05 | 8.97 |
| 2 | 4.94 | 10.35 | 10.11 | 8.33 | 6 | 4.97 | 5.97 | 3.22 | 6.80 | 10 | 12.31 | 7.06 | 4.15 | 3.24 |
| 3 | 8.76 | 15.21 | 4.15 | 5.72 | 7 | 7.72 | 13.15 | 1.29 | 4.66 | 11 | 8.80 | 27.50 | 6.39 | 7.05 |
| 4 | 9.85 | 15.10 | 7.81 | 15.67 | 8 | 3.70 | 9.23 | 5.90 | 3.06 | 12 | 6.91 | 15.93 | 6.48 | 12.30 |

| 96-well Plate Intra-Plate CV (%) for Clonal Growth Outcomes (SFM) |  |  |  |  |  |  |  |  |  |  |  |  |  |  |
| --- | --- | --- | --- | --- | --- | --- | --- | --- | --- | --- | --- | --- | --- | --- |
| T98G ($n = 16$)* | | | | | U251 ($n = 24$)* | | | | | U87 ($n = 16$)* | | | | |
| PID | NC | SP | LE | $\geq 8$ | PID | NC | SP | LE | $\geq 8$ | PID | NC | SP | LE | $\geq 8$ |
| 1 | 10.93 | 8.47 | 10.37 | 24.05 | 1 | 9.59 | 13.15 | 12.26 | 14.55 | 1 | 7.13 | 17.33 | 13.98 | 17.35 |
| 2 | 5.46 | 15.73 | 16.15 | 17.45 | 2 | 13.55 | 8.16 | 10.46 | 9.09 | 2 | 7.48 | 19.56 | 11.30 | 21.03 |

PID - Plate ID; SFM - single-founder microwells; NC - no cells (0 cells); SP - single-cell persistence (1 cell); LE - limited expansion (2-7 cells); $\geq 8$ - $\geq 8$ cells. \* For the 96-well format, cell lines were seeded in separate regions of the same plate; outer wells excluded; $n$ indicates the number of wells (technical replicates) for the cell line.

## 3. Results and Discussion

To improve the scalability and quantitative resolution of proliferative outcomes of clonogenic assays, we developed and evaluated a high-density microwell-based platform enabling large-scale clonal sampling within standard multiwell plate formats. Each well contains approximately 10,000 uniformly sized 50 µm × 50 µm × 50 µm microwells, implemented in both 4-well and 96-well configurations. This architecture enables structured spatial indexing of individual cells and supports longitudinal tracking from seeding through endpoint growth. The cyclic olefin polymer (COP) microwell substrate is coated with PEG to reduce cell adhesion and lateral spreading, thereby promoting confinement of cells within defined microwell boundaries.[18, 19]

To evaluate assay performance across biologically distinct growth behaviors, three widely used glioblastoma (GBM) cell lines with established differences in clonogenic capacity—U251, U87MG, and T98G—were examined using replicate 4-well and 96-well plate formats as described in Section 2.6. Both formats were used to quantify intra-plate uniformity and assess scalability within standard multiwell footprints. The 4-well format additionally allowed comparison across independently prepared plates.

### 3.1 Assay Performance, Seeding Uniformity, and Single-Cell Loading

To evaluate assay performance at the seeding stage, we first quantified the percentage of dispensed cells that entered the microwell compartments and their distribution across individual microwells.

Cells were dispensed into each well at a defined concentration. Following passive sedimentation and orbital shaking, cells settled into microwells. Day 1 label-free brightfield imaging was used to identify and index every microwell within each well. Microwells were classified as either empty or occupied, and the number of cells present in each occupied microwell was determined using the segmentation pipeline described in Section 2.5.

Seeding efficiency was defined as the percentage of dispensed cells that were detected within microwells relative to the total number of cells added to the well on Day 1. Seeding efficiencies averaged across wells from both plate formats were 76% ± 12% for U251, 70% ± 16% for U87MG, and 77% ± 12% for T98G.

To evaluate whether cell entry into microwells followed stochastic loading behavior under passive sedimentation, the distribution of cell counts per microwell was fit to a Poisson model of random occupancy. When all microwells, including empty microwells, were considered, the Poisson mean (λ) values averaged across wells from both 4-well and 96-well plates were 0.88 ± 0.13 for U251, 0.64 ± 0.14 for U87MG, and 0.72 ± 0.12 for T98G. These λ values below 1 indicate sub-saturating loading conditions, under which a substantial fraction of microwells is expected to remain empty and single-cell occupancy is predicted to be the most frequent non-zero outcome.

When the analysis was restricted to occupied microwells only, λ increased to 1.88 ± 0.24 for U251, 1.74 ± 0.13 for U87MG, and 1.65 ± 0.20 for T98G. These values indicate that, among occupied microwells, the average cell count was between one and two cells, with single- and double-cell occupancies accounting for the majority of populated microwells.

These seeding distributions are summarized visually in Figure 4. Stacked bar plots show the mean percentage of occupied microwells classified as single-founder or multi-founder for each plate, averaged across wells. The error bar at the category boundary represents the standard deviation across wells within each plate, reflecting intra-plate variability rather than inter-plate variability. Across both 4-well and 96-well formats, single-founder microwells comprised approximately half of occupied compartments, with the remaining fraction representing multi-founder occupancies. The corresponding mean number of occupied microwells per well is provided below the plots to contextualize sampling depth for each plate.

**Figure 4.**
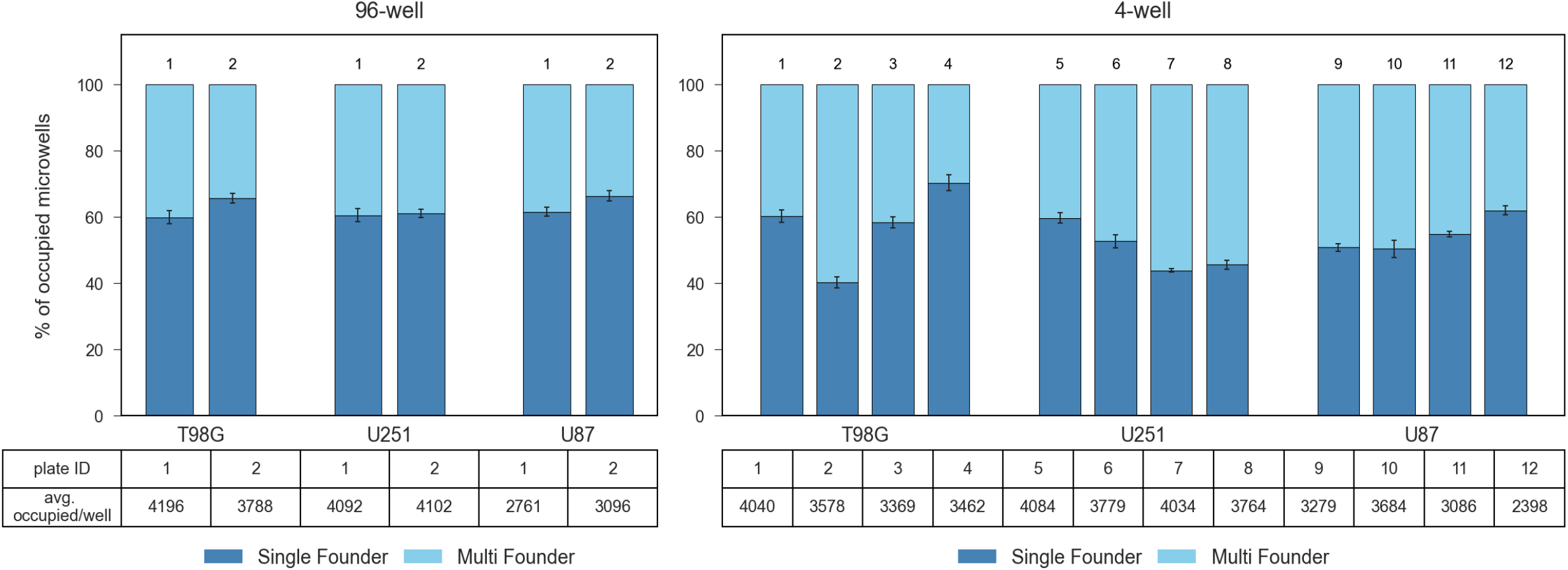
Day 1 founder occupancy distributions for three glioma cell lines (T98G, U251, and U87MG). Stacked bar plots show the mean percentage of occupied microwells classified as single-founder or multi-founder for each plate, averaged across wells within that plate. The error bar at the category boundary represents the standard deviation of the single-founder fraction across wells within the same plate (intra-plate variability). Individual plates are labeled numerically (1-12) above each bar. Table below the plot reports the corresponding mean number of occupied microwells per well for each plate. Data are shown for two 96-well microwell array plates (left) and twelve 4-well microwell array plates (right).

Intra-plate reproducibility for occupancy was evaluated by calculating the coefficient of variation (CV) for the fraction of occupied microwells in each well. CV values for occupancy ranged from 0.7-5.2% for the 4-well format and 5.5-7.9% in the 96-well format. Intra-plate reproducibility of founder distributions was evaluated using CV for the fraction of microwells classified as single-founder and multi-founder within each plate. In the 4-well format, CV values ranged from 2.0-6.5% for single-founder microwells and 1.0-8.8% for multi-founder microwells. In the 96-well format, CV values ranged from approximately 4.3-8.5% for single-founder microwells and 6.7-11.1% for multi-founder microwells (Table 1). These low within-plate CV values indicate consistent seeding behavior across technical replicates in both formats. The combination of stochastic loading behavior, low intra-plate variability, and stable capture efficiency supports the use of this microwell configuration as a quantitative starting point for downstream clonal growth outcome analysis.

### 3.2 Quantitative Distribution of Single-Founder Clonal Growth Outcomes

Single-founder microwells were evaluated from Day 1 classification through Day 6 outcomes using the predefined operational categories described in Section 2.4. Briefly, outcomes were assigned based on Day 6 cell count within the indexed microwell as: 0 cells (no cells; NC), 1 cell (single-cell persistence; SP), 2-7 cells (limited expansion; LE), or ≥8 cells (Figure 5, Table 2).

**Figure 5.**
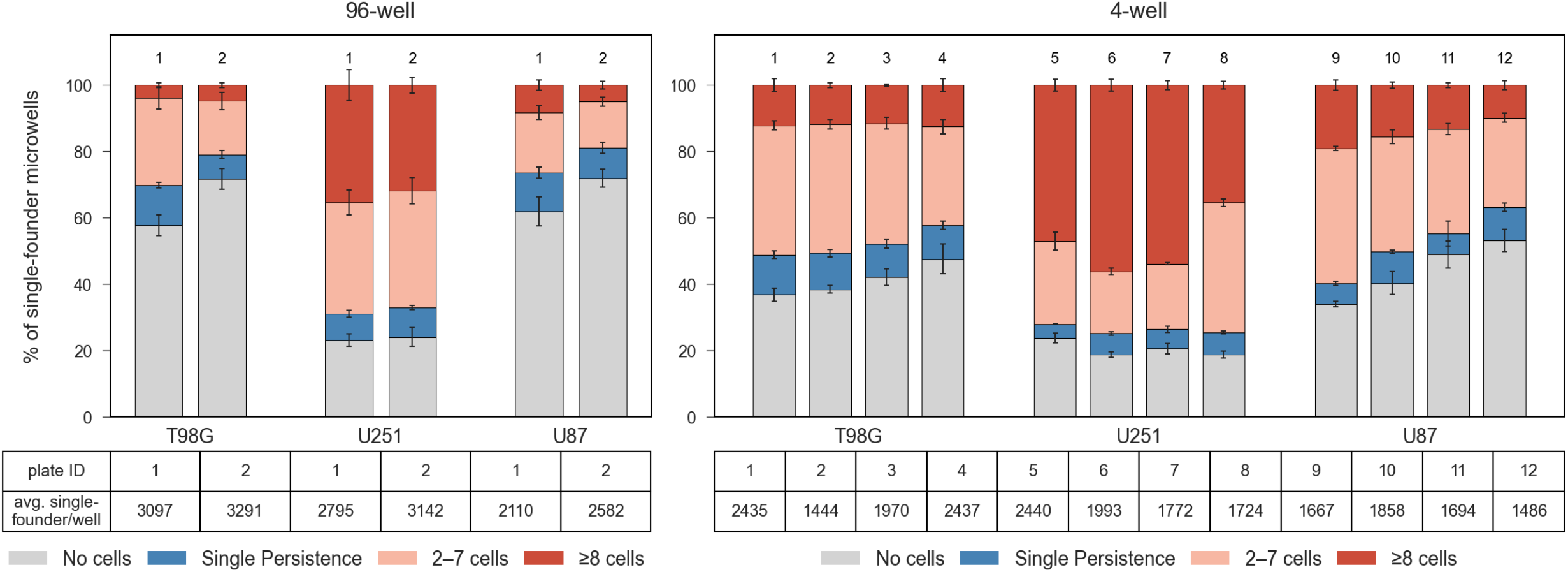
Day 6 clonal growth outcome distributions of single-founder microwells in three GBM cell lines (T98G, U251, and U87MG). Stacked bar plots show the mean percentage of single-founder microwells per well classified into four Day 6 cell count categories: No cells (NC, 0 cells); single-cell persistence (SP, 1 cell); limited expansion (LE, 2-7 cells); ≥8 cell expansion. Values are averaged across wells within each plate. The error bar at the category boundary represents the standard deviation of the outcome fraction across wells within the same plate (intra-plate variability). Individual plates are labeled numerically above each bar. Data are shown for two 96-well microwell array plates (left) and twelve 4-well microwell array plates (right).

These outcome distributions are summarized visually in Figure 5. For each plate, stacked bars show the mean percentage of single-founder microwells in each outcome category, averaged across wells. The error bar at the category boundary represents the standard deviation across wells within the same plate, reflecting intra-plate variability. Across both the 4-well and 96-well formats, distinct distribution patterns were observed among the three GBM cell lines. U251 showed a larger fraction of microwells in the ≥8-cell category, whereas U87MG and T98G displayed higher proportions of NC and SP outcomes. Limited-expansion (2-7 cells) outcomes were observed across all lines but comprised a greater fraction of the total in U87MG and T98G relative to U251. The overall distribution patterns were consistent across plates within each plate format.

Across both plate formats, intra-plate reproducibility of outcome distributions was high. In the 4-well format, intra-plate CVs ranged from 7-28% for SP, 1-16% for LE, and 3-18% for ≥8-cell outcomes, depending on the cell line. The 96-well format showed comparable intra-plate consistency, with CVs of 8-20% for SP, 10-16% for LE, and 17-24% for ≥8-cell outcomes across the three lines (Table 2).

Distinct outcome distributions were consistent across plate formats. U251 exhibited the highest proportion of microwells containing ≥8 cells by Day 6, with approximately one-third to nearly half of single-founder microwells reaching this range. In contrast, U87MG and T98G were characterized by a greater proportion of microwells remaining at low cell counts, with approximately half of single-founder microwells containing either NC or SP, and fewer than 15% reaching ≥8 cells.

These distribution patterns are consistent with published reports describing U251 as strongly clonogenic and U87MG and T98G as comparatively weakly clonogenic lines.[4, 5, 14, 30-34] The agreement with established clonogenic hierarchies indicates that the microwell platform captures expected differences in clonal growth potential while providing expanded distribution-level resolution beyond conventional endpoint colony counts.

A key feature of the assay is its ability to resolve expansion states characterized by minimal or absent expansion, which are not distinguished in conventional colony-formation readouts. Across all three cell lines, a subset of single-founder microwells contained exactly one cell at Day 6 (Fig. 5). These single-cell persistence outcomes were observed more frequently in U87MG and T98G and less frequently in U251.

The ability to quantify single-cell persistence and limited expansion tiers extends the dynamic range of measuring clonal growth outcomes beyond binary colony scoring. By maintaining graded expansion potential, the microwell framework preserves distribution-level information across low-, intermediate-, and high-expansion outcomes. Glioblastoma models are known to exhibit heterogeneous growth behaviors, which is the result of variable proliferative capacities and diversified genetic architecture.[26, 27, 37-39] While the present study does not investigate the molecular basis of these endpoint states, the microwell platform enables systematic quantification of low- and intermediate-expansion outcomes that are not typically distinguished in standard clonogenic workflows.

### 3.3 Multi-Founder Clonal Growth Outcomes

Multi-founder microwells (>1 cell at Day 1) were evaluated as a separate founder class using the same predefined outcome categories described in Section 2.4. At Day 6, outcomes were assigned based on the number of cells present within each indexed microwell as: 0 cells (NC), 1 cell (reduction to single cell; RS), 2-7 cells (limited expansion; LE), or ≥8 cells (Fig. S3). These distributions are summarized in Supplementary Figure S3. For each plate, stacked bars show the mean percentage of multi-founder microwells in each outcome category, averaged across wells. The error bar at the category boundary represents the standard deviation across wells within the same plate, indicating intra-plate variability.

Across both the 4-well and 96-well formats, distinct distribution patterns were observed among the three GBM cell lines. U251 showed a larger fraction of ≥8-cell outcomes, whereas U87MG and T98G exhibited higher proportions of NC, RS, and LE outcomes. The relative ordering of growth outcome distributions across the three lines was consistent with that observed for single-founder microwells. Intra-plate reproducibility of multi-founder outcome distributions was comparable to that observed for single-founder microwells (Table S1).

## 3. Conclusion

In conclusion, we present a high-density microwell-based platform for quantitative assessment of clonal expansion within standard multiwell plate formats. By consolidating large numbers of clonal units within a single plate footprint, the microwell configuration supports high-sampling quantitative analysis using standard automated imaging systems. Importantly, the predefined outcome framework links the initial cell count in each microwell to its clonal growth outcome, allowing single-cell persistence, limited expansion, and higher cell-count outcomes to be quantified separately rather than collapsed into a single non-colony category.

Across seeding and proliferation measurements, the assay demonstrated consistent intra-plate repeatability of founder distributions and linked clonal growth outcomes in both 4-well and 96-well configurations. This structure preserves distribution-level information while maintaining analytical clarity and expands the analytical resolution of clonogenic assessment within conventional multiwell workflows.

There remains an opportunity to further improve assay performance. In future iterations of the platform, single cell microwell occupancy and the coefficient of variation (CV) across technical and biological replicates can be further optimized. Implementation of a lateral flow-based cell seeding strategy may improve single-cell microwell occupancy, while optimization of PLL-g-PEG surface coating may further enhance both single-cell and multi-cell proliferation outcomes.

## Author Contributions

KS, SR, and DPD developed and implemented the assay workflow. BP and DPD designed and executed the microwell plate fabrication process. KS performed the experiments, and SR carried out the data acquisition. SR developed the computational analysis workflow and performed the quantitative analysis of the results. KS, SR, and DPD interpreted the results and jointly prepared the manuscript.

## Data Availability and Code Accessibility

Data and code will be made available upon request.

## Declaration of generative AI and AI-assisted technologies in the manuscript preparation process

During the preparation of this work the author(s) used ChatGPT in order to improve the clarity of the content being delivered and for final proof reading. After using this tool/service, the author(s) reviewed and edited the content as needed and take(s) full responsibility for the content of the published article.

## Acknowledgements

We gratefully acknowledge funding support from the Cancer Prevention & Research Institute of Texas (CPRIT), project DP240117.

