## Supplement for "A Scalable High-Density Microwell Assay for Single-Cell Clonal Expansion Profiling"

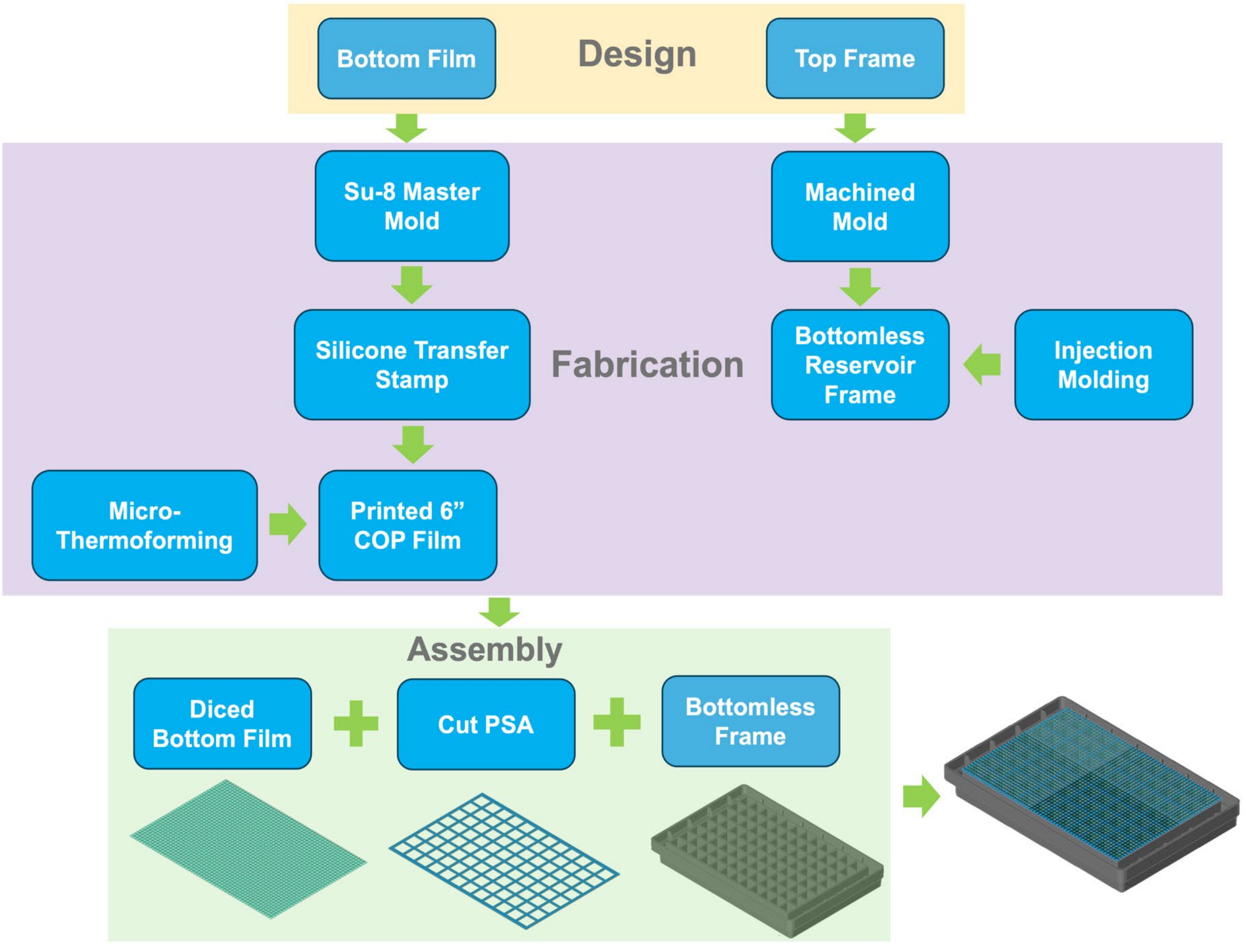


Figure S1. Schematic workflow of the microwell assay plate manufacturing process. The process begins with the design of two components: a microwell array bottom film and a bottomless top frame. Fabrication of the bottom film involves photolithographic generation of an SU-8 master mold, followed by replication of a silicone negative transfer mold. This mold is used to pattern microwell features onto a 6-inch cyclic olefin polymer (COP) film via a micro-thermoforming process. The top frame is manufactured separately using a conventional injection molding process. The two components are subsequently aligned and assembled by lamination using a die-cut biocompatible pressure-sensitive adhesive (PSA), forming the final microwell assay plate.

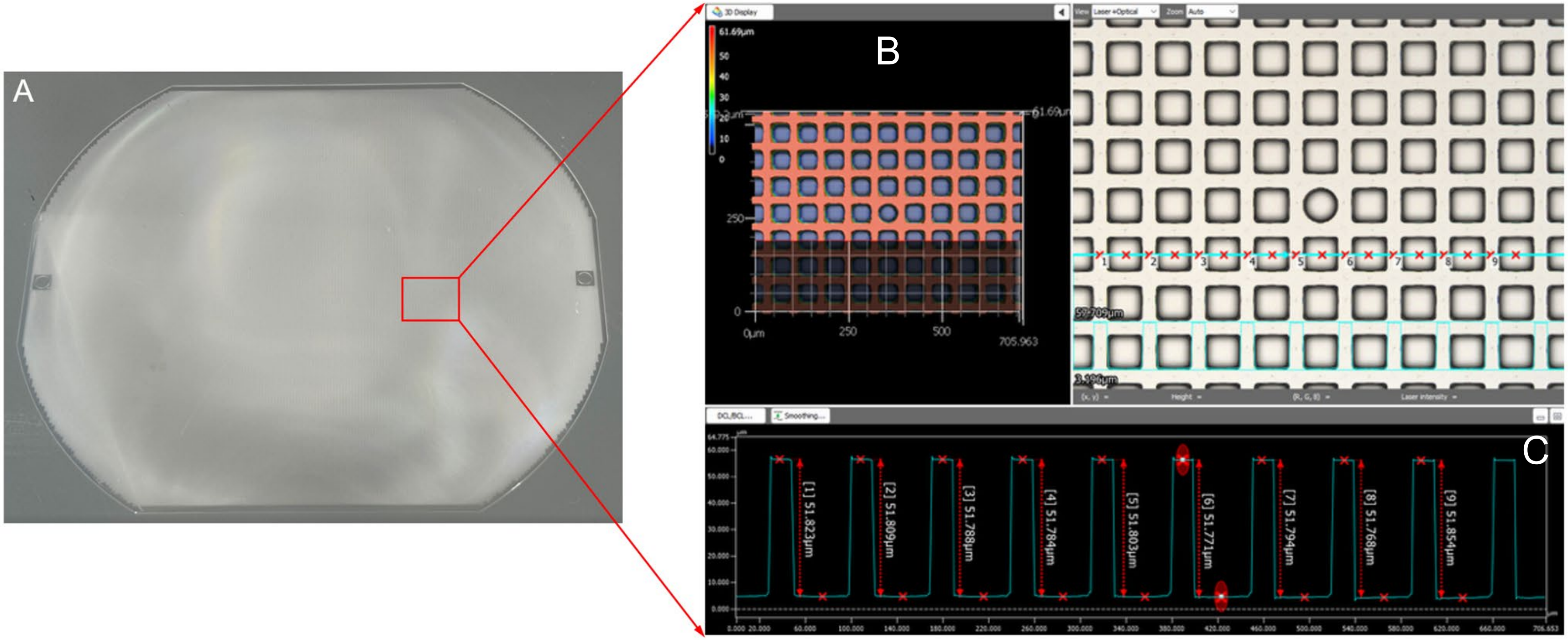


Figure S2. Thermoformed COP film and dimensional metrology. (A) Array of microwells thermoformed into a 6-inch COP film (188 µm thickness). Individual pieces were subsequently diced to generate 4-well and 96-well plate formats. (B) Representative microscopic image of a randomly selected region used to quantify microwell dimensions. (C) Profilometer-derived depth profiles across a row of microwells. Dimensional variability was <0.1%, indicating excellent feature uniformity and reproducibility.

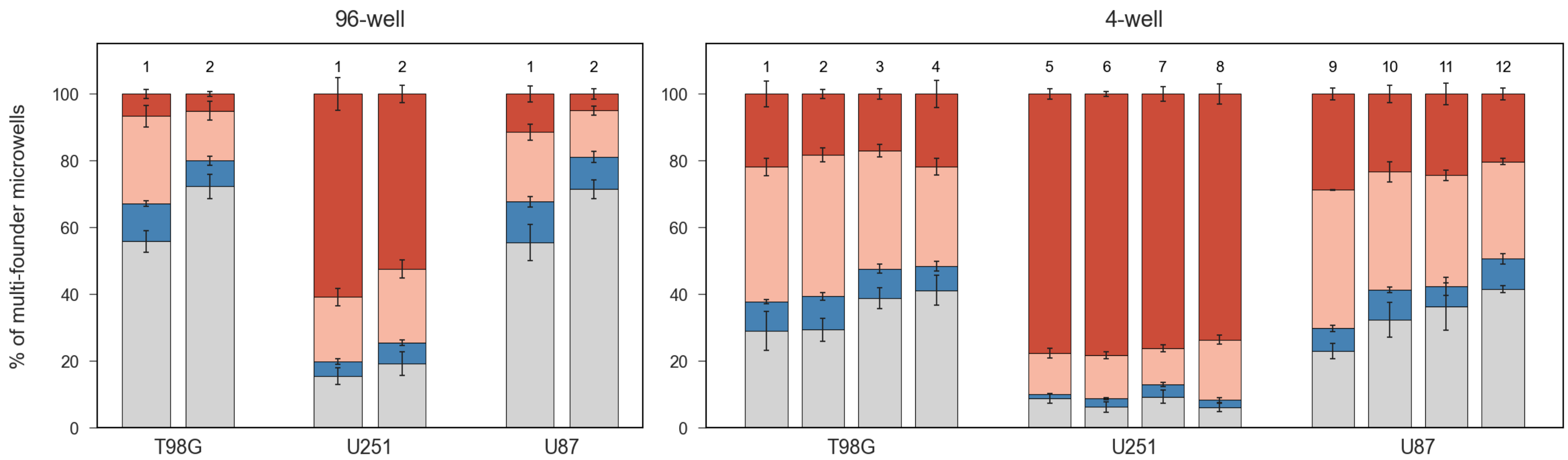


| plate ID | 1 | 2 | 1 | 2 | 1 | 2 |
|---|---|---|---|---|---|---|
| avg. multi-founder/well | 2107 | 1728 | 1817 | 1997 | 1316 | 1307 |

| 1 | 2 | 3 | 4 | 5 | 6 | 7 | 8 | 9 | 10 | 11 | 12 |
|---|---|---|---|---|---|---|---|---|---|---|---|
| 1604 | 2134 | 1399 | 1026 | 1644 | 1786 | 2262 | 2040 | 1612 | 1825 | 1391 | 912 |



Figure S3. Day 6 clonal growth outcomes of multi-founder microwells in three GBM cell lines (T98G, U251, and U87). Histograms show the mean percentage and number of multi-founder microwells per well exhibiting one of four outcomes between day 1 and day 6: No cells (NC, 0 cells); reduction to a single (RS, 1 cell); limited expansion (LE, 2-7 cells); ≥8 cell expansion. Data are shown for two 96-well microwell array plates (left) and twelve 4-well microwell array plates (right). Reported values represent the mean number of microwells per well (i.e., averaged across wells within each device).

| 4-well Plate Intra-Plate CV (%) for Clonal Growth Outcomes (MFM) | | | | | | | | | | | | | | |
|---|---|---|---|---|---|---|---|---|---|---|---|---|---|---|
| T98G ($n = 4$) | | | | | U251 ($n = 4$) | | | | | U87 ($n = 4$) | | | | |
| PID | NC | RS | LE | ≥8 | PID | NC | RS | LE | ≥8 | PID | NC | RS | LE | ≥8 |
| 1 | 21.95 | 7.72 | 5.07 | 18.06 | 5 | 13.45 | 12.80 | 17.01 | 6.14 | 9 | 8.51 | 15.63 | 4.19 | 8.39 |
| 2 | 9.81 | 10.74 | 9.63 | 9.09 | 6 | 23.30 | 16.65 | 10.69 | 4.61 | 10 | 17.64 | 15.90 | 10.92 | 10.76 |
| 3 | 7.25 | 19.37 | 6.14 | 13.77 | 7 | 20.55 | 17.11 | 8.74 | 3.45 | 11 | 24.41 | 31.10 | 4.49 | 13.21 |
| 4 | 13.72 | 13.30 | 9.60 | 24.92 | 8 | 17.01 | 31.46 | 1.59 | 8.92 | 12 | 9.61 | 17.73 | 8.74 | 10.67 |
| 96-well Plate Intra-Plate CV (%) for Clonal Growth Outcomes (MFM) | | | | | | | | | | | | | | |
| T98G ($n = 16$)* | | | | | U251 ($n = 24$)* | | | | | U87 ($n = 16$)* | | | | |
| PID | NC | RS | LE | ≥8 | PID | NC | RS | LE | ≥8 | PID | NC | RS | LE | ≥8 |
| 1 | 22.13 | 10.49 | 10.50 | 14.22 | 1 | 11.32 | 16.00 | 15.53 | 19.51 | 1 | 11.32 | 16.00 | 15.53 | 19.51 |
| 2 | 21.40 | 9.51 | 21.40 | 20.98 | 2 | 10.66 | 19.03 | 12.95 | 32.61 | 2 | 10.66 | 19.03 | 12.95 | 32.61 |

PID - Plate ID; MFM - multi-founder microwells; NC - no cells (0 cells); RS - reduced to single cell (1 cell); LE - limited expansion (2-7 cells); ≥8 - ≥8 cells.

* For the 96-well format, cell lines were seeded in separate regions of the same plate; outer wells excluded; $n$ indicates the number of wells (technical replicates) for the cell line.

**Table S1**
Intra-plate coefficient of variation (CV, %) for Day 6 clonal growth outcomes for multi-founder microwells across 4-well and 96-well formats. Each value represents the intra-plate CV calculated across $n$ wells in a plate.

| 4-well Plate Inter-Plate CV (%) for Seeding | | | | | | | | |
|---|---|---|---|---|---|---|---|---|
| T98G ($n = 4$) | | | U251 ($n = 4$) | | | U87 ($n = 4$) | | |
| Occ. | SFM | MFM | Occ. | SFM | MFM | Occ. | SFM | MFM |
| 8.2 | 22.8 | 30.0 | 4.2 | 16.4 | 14.1 | 17.3 | 9.1 | 27.2 |

Occ. - microwell occupancy; SFM - single-founder microwells; MFM - multi-founder microwells. $n$ indicates the number of plates (biological replicates).

**Table S2**
Inter-plate coefficient of variation (CV, %) for microwell occupancy and founder distribution among occupied microwells. CV was calculated across independent plates using the average value from each plate.

| 4-well Plate Inter-Plate CV (%) for Proliferation Outcomes (SFM) | | | | | | | | | | | |
|---|---|---|---|---|---|---|---|---|---|---|---|
| T98G ($n = 4$) | | | | U251 ($n = 4$) | | | | U87 ($n = 4$) | | | |
| NC | SP | LE | ≥8 | NC | SP | LE | ≥8 | NC | SP | LE | ≥8 |
| 28.9 | 26.3 | 21.6 | 25.4 | 28.2 | 10.5 | 33.1 | 26.0 | 15.7 | 25.8 | 22.0 | 31.4 |

SFM - single-founder microwells; NC - no cells (0 cells); SP - single-cell persistence (1 cell); LE - limited expansion (2-7 cells); ≥8 - ≥8 cells. $n$ indicates the number of plates (biological replicates).

**Table S3**
Inter-plate coefficient of variation (CV, %) for Day 6 clonal growth outcomes in single-founder microwells across replicate 4-well plates. CV was calculated across independent plates using the average value from each plate.
